# Error-driven representation learning in the mesolimbic system

**DOI:** 10.64898/2026.05.18.725950

**Authors:** George Cai, Max F. Scheller, Wolfgang Kelsch, Samuel J. Gershman

## Abstract

In reinforcement learning, an agent learns to map representations of the environment state to predictions of future reward. Most prior work in neuroscience has assumed a fixed representation and studied how reward prediction errors (thought to be conveyed by phasic dopamine signals) are used to update the mapping from representations to predictions. However, work in machine learning has demonstrated that much more powerful predictive systems can be learned by using the errors to update the representations themselves. We study whether the brain does something similar by leveraging simultaneous recordings of striatal projection neurons in the olfactory tubercle (putatively representing state features) and dopamine neurons in the ventral tegmental area. We show that trial-by-trial changes in striatal activity are more consistent with dopamine-driven representation learning than a variety of alternative updating schemes. This result suggests a convergence of representation learning principles in biological and artificial systems.

## Introduction

Learning to predict future reward (the reinforcement learning problem) is a challenge faced by virtually all animals. The brain uses multiple learning algorithms to solve this problem (***Kool et al., 2018***), including error-driven updating of reward predictions. In the simplest version of this idea, the brain caches a look-up table of reward predictions for each state of the environment, increasing the predictions when they underestimate reward (positive prediction error), and decreasing them when they overestimate reward (negative prediction error). While this algorithm will, with enough experience, arrive at correct reward predictions, it is hopelessly inefficient in environments with many states. Multidimensional environments can have a number of states that is exponential in the number of dimensions, making it impossible for animals to gather enough experience in each state—the *curse of dimensionality*. For example, mice have over 1000 olfactory receptor types, yielding an astronomical number of distinct states; caching algorithms cannot succeed in such a vast state space.

A more promising strategy is to learn mappings from some feature space to reward predictions. A classic version of this idea is the *linear function approximator*, where the learned parameters are weights that specify the contribution of each dimension (represented as an input “feature”) to future reward. Linear function approximation has been applied extensively to both animal (***Rescorla and Wagner, 1972; Ludvig et al., 2012***) and machine (***Geramifard et al., 2013***) learning. However, it is fundamentally limited in the kinds of reward predictions it can learn. For example, it cannot learn conjunctions of features, unless the conjunctions are added to the feature set, leading to a combinatorial explosion—the curse of dimensionality again.

A breakthrough in modern machine learning was the development of deep neural networks that can learn to represent complex non-linear functions (***LeCun et al., 2015***). Instead of assuming a fixed feature space, these networks learn to represent features that optimize an objective function (e.g., reward prediction accuracy), using error backpropagation to update all the parameters of the function approximator. This enabled artificial agents to play complex video games that were previously impossible using linear function approximation (***Mnih et al., 2015***). It has been suggested that the brain may similarly use deep representation learning to solve complex tasks (***Richards et al., 2019; Botvinick et al., 2020; Gershman and Ölveczky, 2020***).

While some representations in the brain might be consistent with those learned by deep neural networks, there is currently little direct evidence that the brain uses error-driven updating to acquire these representations (for some indirect evidence and models, see ***Alexander and Gersh- man, 2021; Jakob et al., 2022***). Our main contribution is to directly test this hypothesis, by analyzing data from an experiment in which several components of the learning system were measured simultaneously (***Scheller et al., 2026***).

Dopamine neurons (DANs) in the ventral tegmental area (VTA) and striatal projection neurons (SPNs) in the olfactory tubercle (OTu) were recorded while mice performed a go/no-go task with olfactory cues. To study representation learning in this setup, we made two critical postulates. First, we postulated that dopamine conveys the prediction error used for updating. While extensive evidence supports this postulate for the updating of reward predictions (***Schultz et al., 1997; Gershman et al., 2024***), it is currently unknown whether dopamine also drives representation learning. Second, we postulated that the OTu SPNs, which receive dense dopamine projections from the VTA as well as from olfactory cortex, represent the features used for reward prediction. This postulate is consistent with the role of OTu SPNs in coding odor identity (***Lee et al., 2024***), valence (***Gadziola et al., 2015; Millman and Murthy, 2020; Martiros et al., 2022; Oettl et al., 2020; Winkelmeier et al., 2022***), and sensory associations (***Wesson and Wilson, 2010***). With these postulates in hand, we could address our primary question: do representations (putatively encoded by OTu SPNs) change in the way predicted by backpropagation of errors (putatively encoded by DANs)?

## Results

### Reinforcement learning framework and hypothetical neural architecture

We model an agent learning to predict reward *r* with a value function approximator *v*_*ϕ*_(**x**), where **x** represents sensory input (which we take to be the environment state), and *ϕ* are learnable parameters. The optimization problem is to choose the parameters that minimize mean squared error *L*(*ϕ*) = 𝔼 [*δ*^2^|*ϕ*], where *δ* = *r* − *v* (**x**) is the *reward prediction error* (RPE), and the expectation is taken with respect to the distribution *p*(**x**, *r*). To learn online, the agent collects samples (**x**, *r*) and updates the parameters by following the stochastic gradient:

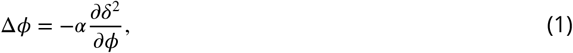

where *α* ∈ [0, 1] is a learning rate. Provided the learning rate decreases over time in accordance with the Robbins-Monro conditions (***Robbins and Monro, 1951***), the value function will asymptotically minimize mean squared error.

We model *v*_*ϕ*_(**x**) as a two-layer linear network with parameters *ϕ* = (*θ*, **w**):

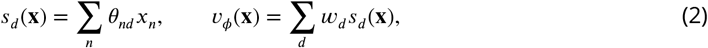

where *n* indexes sensory features and *d* indexes state representation features. For completeness and clarity, the matrix form is

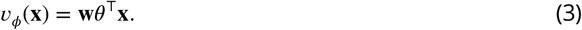

Here, each row of the matrix *θ* is the SPN population response to one of the odor stimuli. Although technically this network is equivalent (in terms of the function class that it can represent) to a one-layer linear network, the gradient descent learning dynamics are distinct due to higher order terms in the weights from the chain rule (***Saxe et al., 2013***). We adopt the linear structure for simplicity; our approach can be readily extended to non-linear networks.

The stochastic gradient updates for the parameters are given by:

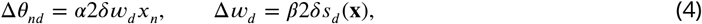

where we have assumed separate learning rates (*α* and *β*) for the two updates. We also assume a one-hot encoding of sensory inputs, where *x*_*n*_ = 1 if stimulus *n* is presented (0 otherwise). In this case, the update for *θ*_*nd*_ is proportional to *δw*_*d*_ whenever stimulus *n* is presented, and otherwise stays unchanged.

We propose a mapping from this abstract framework to a neural circuit architecture (Figure 1**A**). For odor stimuli, we assume that sensory inputs **x** are encoded in the olfactory bulb and piriform cortex. In general, the encoding is not one-hot, but this assumption captures the fact that highly discriminable odors are encoded as distinct “odor objects” (***Wilson and Sullivan, 2011***), such that odor identity can be decoded with high accuracy using a sparse linear readout (***Miura et al., 2012; Mathis et al., 2016***). The olfactory areas send outputs to the OTu, which also receives dopamine input from the VTA. We assume that SPNs in the OTu encode the state representation **s**, while VTA dopamine neurons encode the RPE *δ*. Over the course of learning, the stimulus representations re-organize so that different value categories become linearly decodable (***Millman and Murthy, 2020; Oettl et al., 2020; Winkelmeier et al., 2022***). The OTu projects to an intermediate neuron layer that comprises ventral pallidum, VTA GABA neurons and other relays that represents the reward expectation—the putative state value *v*. Lastly, the value information is transmitted to VTA DANs to compute the RPE.

**Figure 1.**
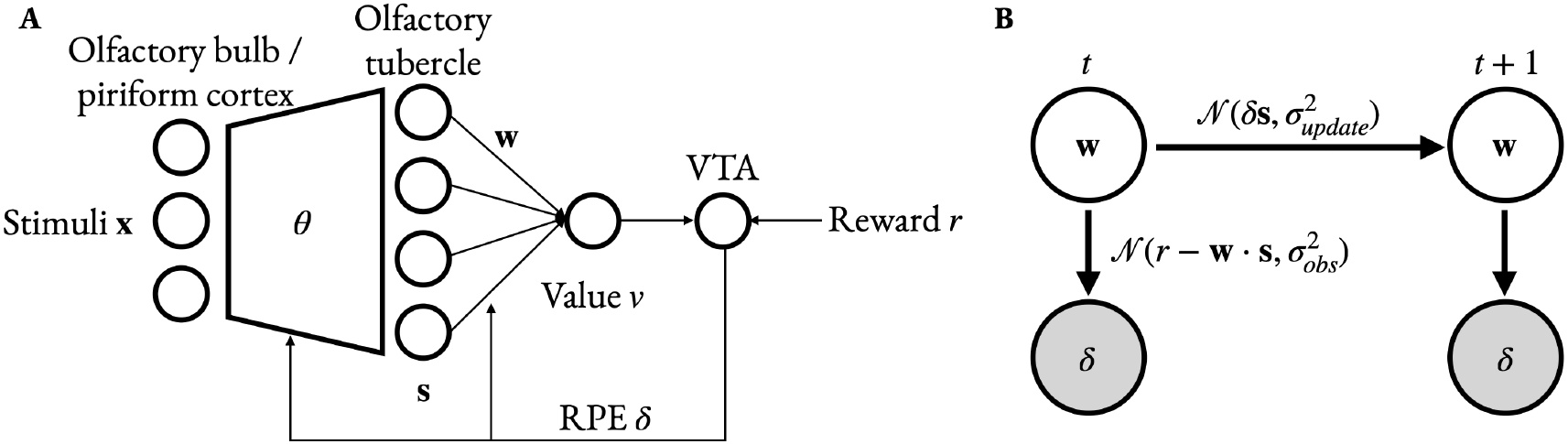
Neural circuit model for learning values of odor-cued states. (**A**) Stimulus information is transmitted to the olfactory tubercle (OTu) of the ventral striatum via the olfactory bulb and/or piriform cortex. Striatal projection neurons (SPNs) in OTu represent states **s**(**x**) = *θ*^⊤^**x**, and a learned weighted sum of SPN activity estimates the value *v*_*ϕ*_(**x**) = **w** ⋅ **s**(**x**). The reward prediction error (RPE) *δ* = *r* − *v*_*ϕ*_(**x**) is projected back from the ventral tegmental area (VTA) to update both feature representation parameters *θ* and the linear approximation parameters **w**.(**B**) Linear-Gaussian dynamical system model of weight updates. Shaded variables are observed while unshaded variables are latent.

To make predictions about representation learning in OTu, we need to understand how the updates in Eq. 4 should manifest in OTu activity changes. Plugging the update into Eq. 2 and applying the one-hot assumption for **x** yields:

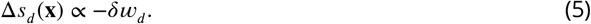

This implies that OTu neuron *d* should increase its activity if the signs of *δ* and *w*_*d*_ match, or decrease its activity if the signs mismatch.

### Probabilistic inference over weight trajectories

Eq. 5 poses an immediate obstacle to testing the model predictions: while we can ostensibly measure *δ* from the activity of DANs, we cannot directly measure *w*_*d*_, which is putatively a synaptic strength. We can, however, infer *w*_*d*_ under certain assumptions.

The linearity of our model allows us to apply Bayesian methods (Rauch-Tung-Striebel smoothing) to solve the weight inference problem, provided we additionally assume Gaussian noise on both the weight updates and DAN measurements (Figure 1B). The technical details can be found in the Methods. The output of this procedure is a trajectory of inferred weights, which we can plug into Eq. 5 to make directional predictions.

Although we cannot experimentally validate the inferred weight trajectories, we show that weight recovery is reliable in synthetically generated SPN and DAN activity (Figure 2**A**). Synthetic model parameters were set to values inferred by likelihood optimization on the corresponding experimental data (see Methods for details). We also evaluated the inferred representation updates in the synthetic setting. To avoid additional assumptions about the initial representation or inferring a learning rate, we computed correlations between the updates Δ*s*_*d*_ (**x**) and *δw*_*d*_. The posterior predictive check shows adequate update recovery in Figure 2**B**. In addition, the update prediction is well-correlated with the ground truth updates in the simulated data in Figure 2**C** (gradient descent rule). Alternative learning rules are discussed in the following section.

**Figure 2.**
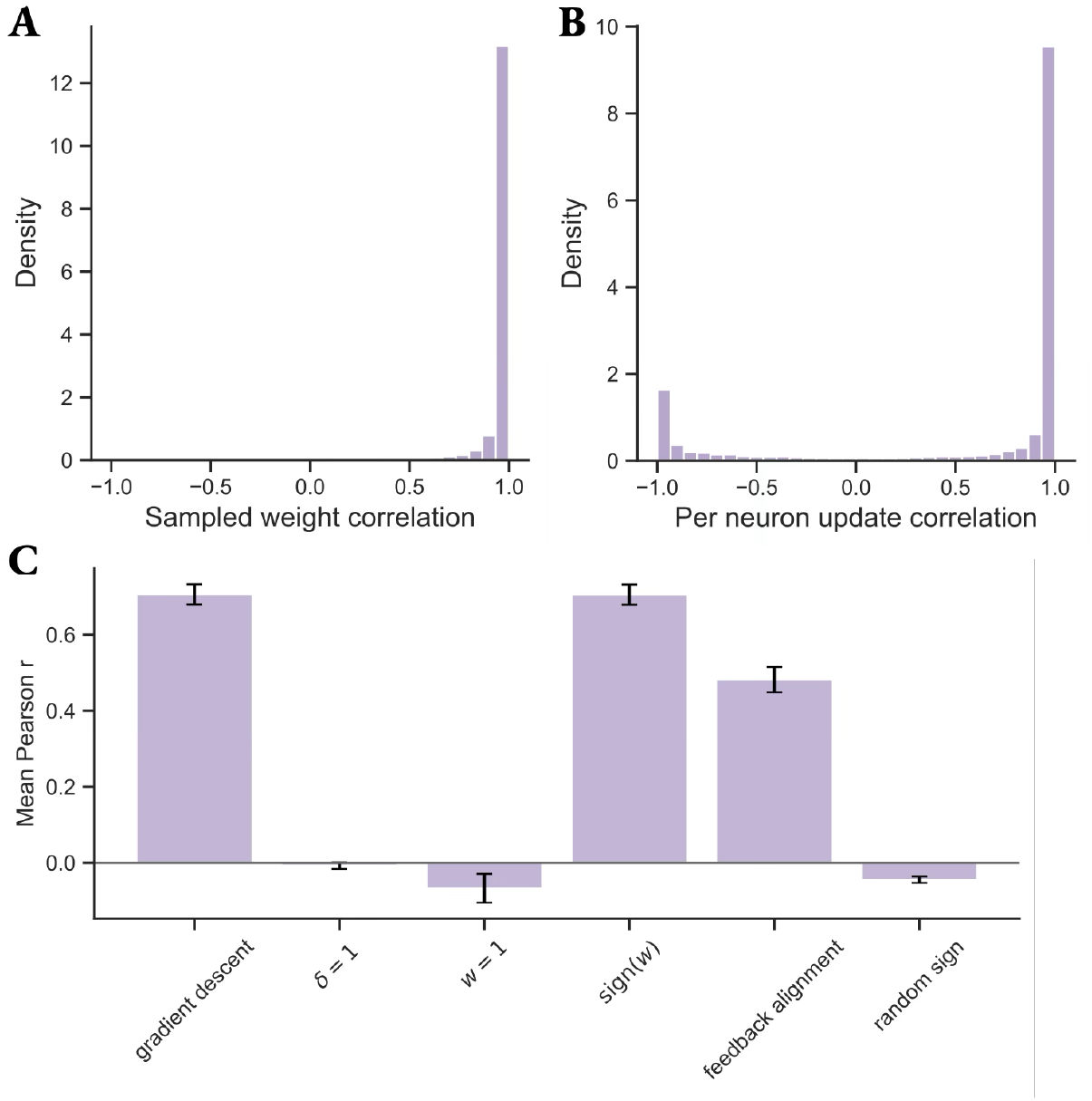
Validation of weight inference on synthetic data. (**A**) Correlation between the true and inferred weight trajectory. (**B**) Correlation between true and reconstructed representation updates from sampling weight trajectories and reconstructing RPEs and representation trajectories. **C** Correlation between true and inferred representation updates using weight trajectory mean and synthetic RPEs. Bars show mean and standard error across neurons.

### Changes in OTu activity are consistent with error-driven representation learning

We now apply our modeling framework to the experimental data collected in the experimental configuration shown in Figure 3**A** (more details in Methods). Head-fixed mice (under mild water restriction on the experimental days) were presented three odors associated with different probabilities of water reward (0%, 50%, 100%) in a trial-based format (Figure 3**B**). After reaching criterion behavioral performance (correct anticipatory licking in the 50% and 100% trials and withholding licking in the 0% trials), the reward probabilities of the 0% and 100% odor cues were reversed. In the reinforcement learning formalism, odors (**x**) cue different states, evoking OTu SPN responses **s**(**x**). At reward time, VTA DAN signal the RPE *δ*. With this mapping, we can combine the DAN activity measurements with the inferred weight trajectory to directly test whether OTu SPN activity follows Eq. 5 on a trial-by-trial basis.

**Figure 3.**
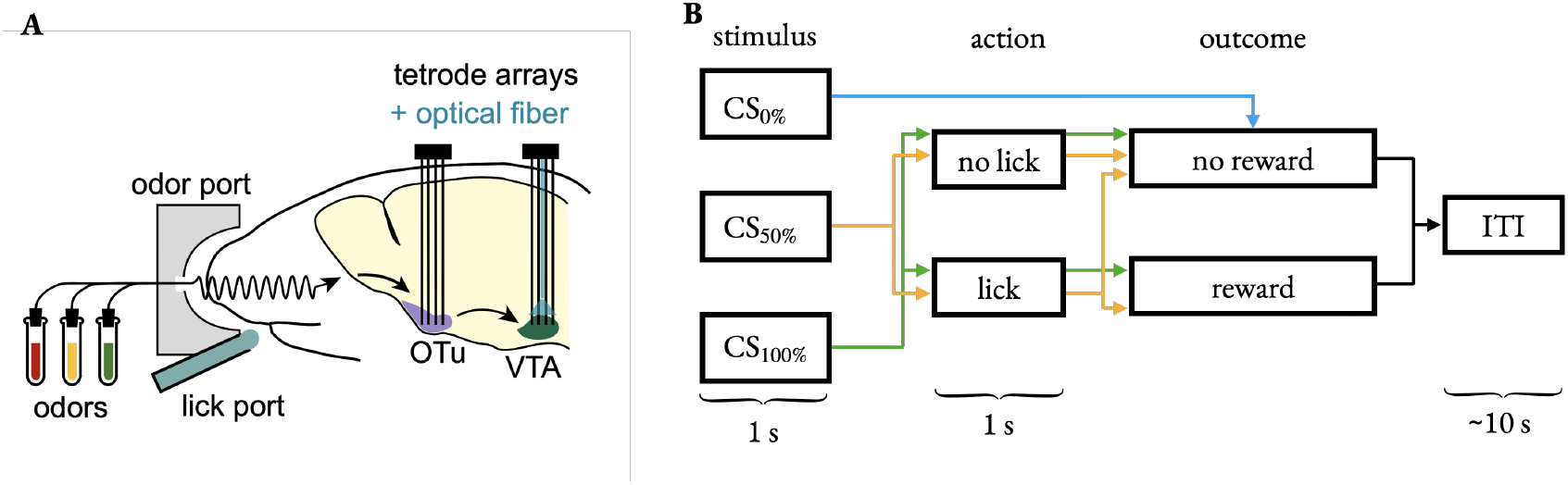
Experimental setup. (**A**) Simultaneous recording in the olfactory tubercle (OTu) and ventral tegmental area (VTA). Licking and neural data were recorded in a head-fixed configuration during stimulus and reward presentation. (**B**) Behavioral paradigm. Mice were trained on a go/no-go task with three odor stimuli that predicted reward with 0%, 50%, and 100% probability. ITI: intertrial interval.

We find that gradient descent produces state representation updates with significantly higher SPN correlations than alternative models (Figure 4; full distributions in Figure S1). We show that *δ* and **w** are individually important for predicting SPN updates by setting *δ* = 1 or **w** = **1**. We note that **w** = **1** can be thought of as assuming SPNs are value-coding since the value estimate becomes a sum over **s**. Keeping only the sign of **w** achieves relatively high correlations, but still below the full gradient.

**Figure 4.**
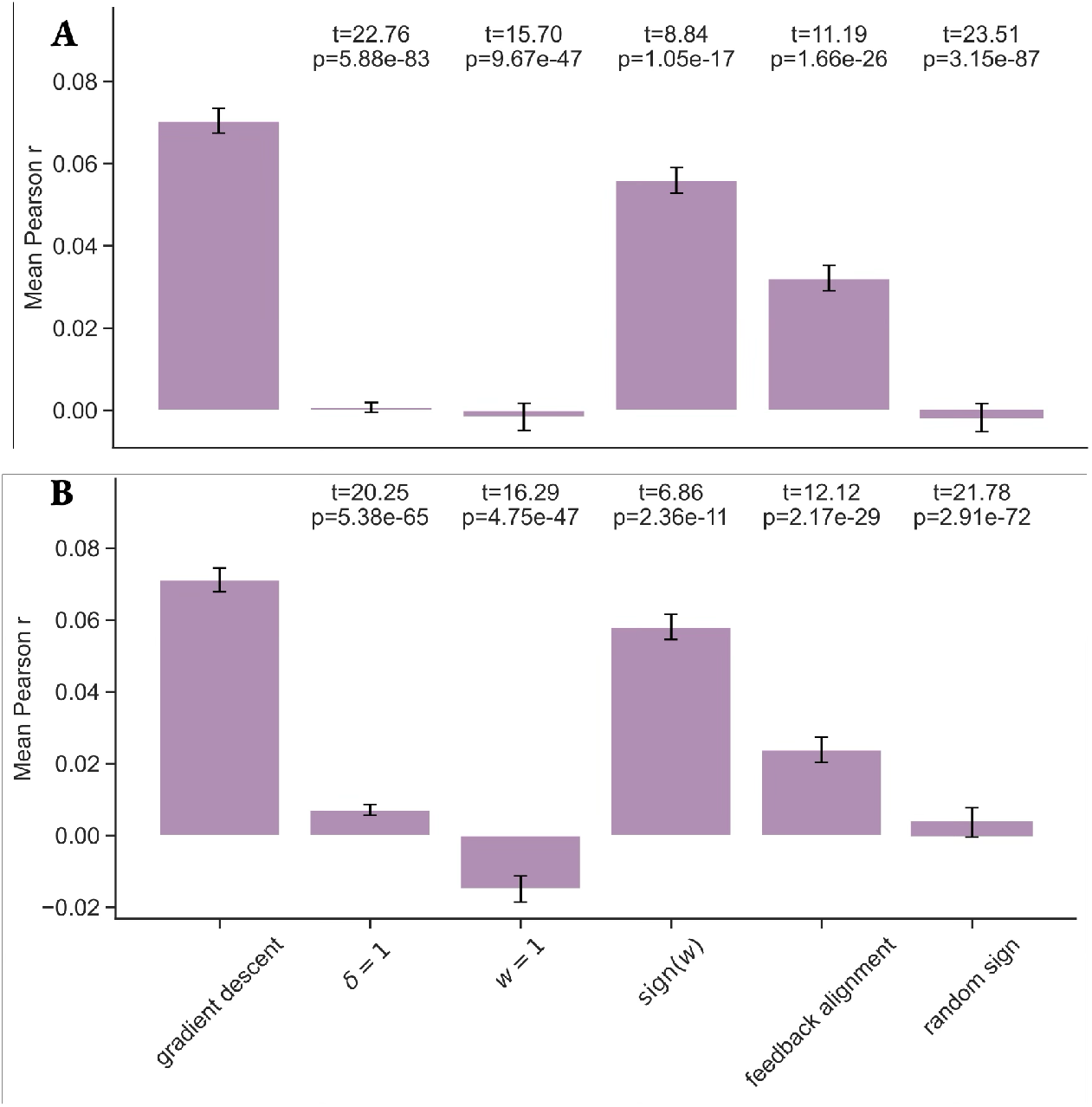
Correlations between predicted and observed changes in olfactory tubercle activity during (**A**) Pavlovian conditioning sessions (411 neurons) and (**B**) reversal sessions (284 neurons). Bars show mean and standard error across neurons. The gradient model (Eq. 5) was compared to several alternative representation updates: fixing *δ* = 1, fixing **w** = 1, replacing **w** by its element-wise sign (sign(**w**)), replacing **w** by a fixed Gaussian vector (feedback alignment), and randomly sampling the sign of the update. The *t*-values and *p*-values shown above each alternative model are derived from paired *t*-tests.

We also test an alternative model of feedback alignment, where **w** is replaced by a random Gaussian vector (***Lillicrap et al., 2016***). This rule is considered more biologically plausible than gradient descent because it avoids the transport of the SPN post-synaptic weights to the pre-synaptic connection, but the representation update correlations are significantly lower than gradient descent (Figure 4). Lastly, we check that update correlations are significant relative to the noise-driven representation changes by randomly sampling the sign of the update. These results are consistent in both Pavlovian conditioning and contingency reversal sessions (0% and 100% reward cues are reversed), which requires updates in opposite directions since the RPE sign flips in early reversal trials. We show the results are robust to pre-CS window choice in Figure S2 and threshold number of neurons recorded in Figure S3.

To show more directly the data underlying these correlations, Figure 5 plots the mean SPN update as a function of the theoretical update, for each of the hypothetical algorithms described above. These plots illustrate that only the stochastic gradient and its sign transform generate accurate predictions.

**Figure 5.**
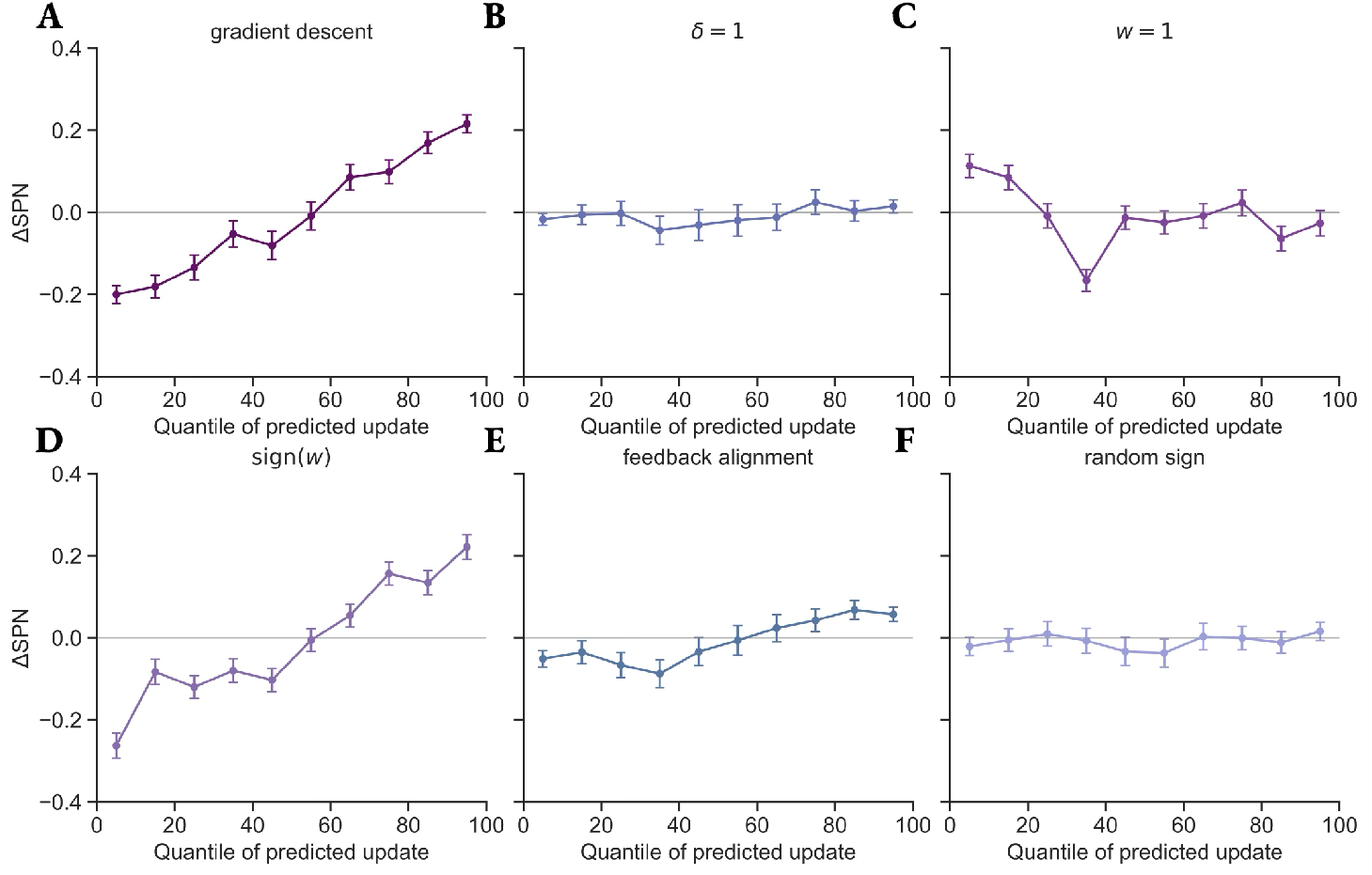
Change in SPN response between successive representations of an odor as a function of the decile of predicted change according to (**A**) gradient descent, (**B**) *δ* = 1, (**C**) **w** = **1**, (**D**) sign(**w**), (**E**) feedback alignment, (**F**) random sign update rules. Points denote mean and error bars denote standard error. A total of *n* = 99902 updates were considered across all SPNs and trials and sessions of both types.

We verify that the greater predictivity of the gradient descent rule cannot be explained by different numbers of hyperparameters for different learning rules in the Kalman smoother weight inference. We use an 80%-20% train-test split to fit hyperparameters on the train split, run weight inference on the full session, and compute update correlations on the test split (see Held out comparison for learning rules for details). We confirm that the full gradient descent rule best predicts SPN response changes on successive trials in Figure S4 and Figure S5. Lastly, we use the Bayesian information criterion (BIC) to compare the alternative weight dynamics models’ ability to explain the data in Figure 6. We find that the gradient descent model has the lowest BIC on 24/26 sessions analyzed.

**Figure 6.**
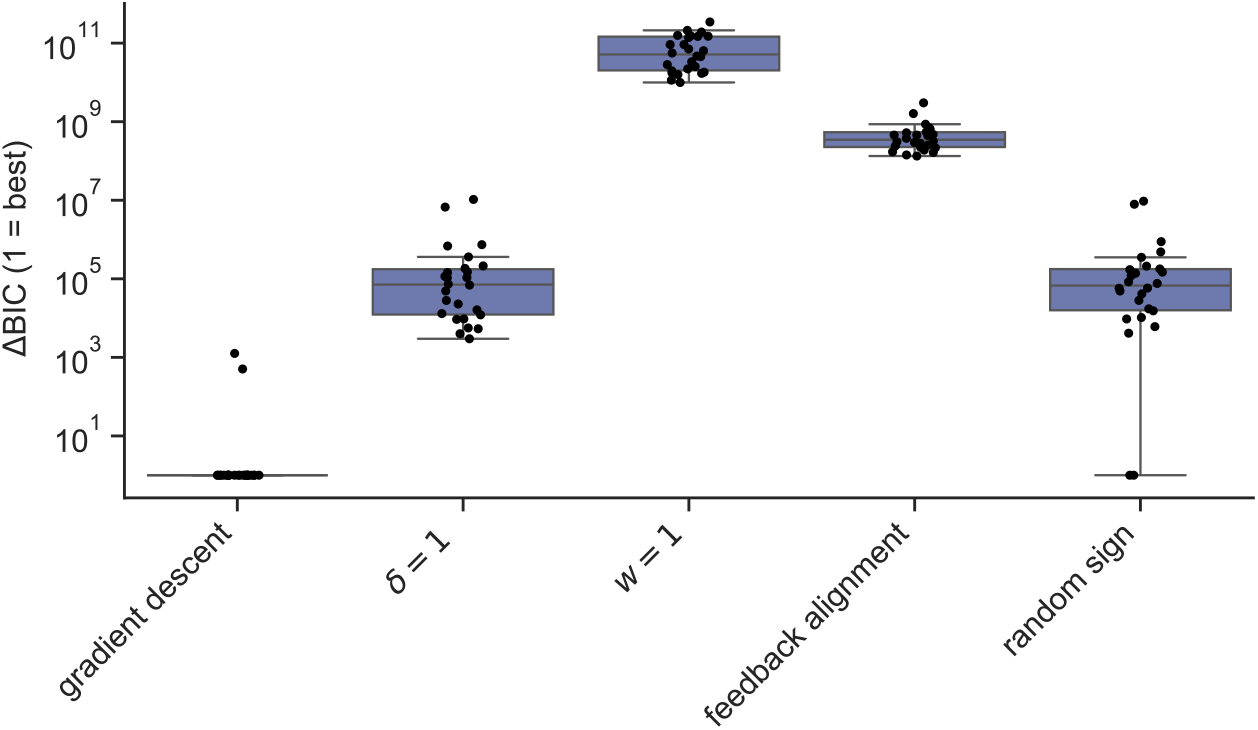
Comparison between alternative weight dynamics models on the ability to explain the SPN and DAN response data using the Bayesian information criterion. Each point is subtracted from the minimum BIC value for that session and offset by 1 for visualization purposes on a logarithmic scale. Box denotes *Q*_1_, *Q*_2_, *Q*_3_, and whiskers denote *Q*_1_ − 1.5IQR, *Q*_3_ + 1.5IQR where *Q*_*i*_ denotes *i*th quartile.

### Validating assumptions and checking robustness

In our model, we made multiple simplifying assumptions. First, using linear feature representations (*s*_*d*_ (**x**) = ∑_*n*_ *θ*_*nd*_ *x*_*n*_) implies that updates to one state’s representation do not influence another state’s representation. We probe the validity of this assumption by computing correlations for intervals between same-stimulus presentations and updates. If there are more intervening trials between successive presentations of a particular stimulus, then we expect larger absolute changes to this stimulus if the representations are coupled. In Figure S6, we find the inter-update interval for a given stimulus has a significant non-zero mean correlation with the per-neuron updates only in the reversal sessions. This result is partially consistent with our model’s assumption that updates to one representation does not affect the representation of another stimulus.

Second, we chose a decaying learning rate (see Methods) to guarantee convergence of stochastic gradient descent to the value function approximation that minimizes mean squared error given fixed features (***Sutton and Barto, 2018***). However, the brain may not modulate the synaptic learning rate this way. We show in Figure S7 that our results are robust to holding *β* constant across trials and inferring it via maximum likelihood. In reality, the learning rate may take an intermediate sequence of values between these extremes.

## Discussion

Much attention has been devoted to neural representations of state suitable for reinforcement learning (e.g., ***Rao, 2010; Stachenfeld et al., 2017; Garvert et al., 2017; Schuck et al., 2016; Hennig et al., 2023***), which are thought to underlie the efficiency and flexibility of animal behavior. However, relatively fewer studies ask how these state representations are developed or learned, especially with grounding in neural data (see ***Niv, 2019***, for a review). Developmental programs, un-supervised learning, and reinforcement learning are all plausible contributors to adapting these neural representations, and it is of interest to identify the underlying computations and algorithms. In this work, we demonstrate that the computation of RPE minimization implemented by the gradient descent algorithm is predictive of trial-by-trial changes in stimulus representations in the OTu. We provide a simple neural circuit model of error-driven representation learning of odor states in the VS-VTA circuit. Next, we will discuss related work on state representations being sculpted by dopamine signals and prediction error minimization.

### Dopamine-based representation learning

Our analysis supports the hypothesis that dopamine provides an error signal for state representation updates in OTu for odor-determined states.

In a similar dual site recording configuration in OTu and VTA, pairing one odor’s presentation to optogenetically induced VTA dopamine release reinforces stimulus representations and increases their separability due to changes in the paired stimulus (***Oettl et al., 2020***). Similarly, in a task with a contingency reversal paradigm with only go/no-go stimuli, OTu SPN reinforced their responses to rewarded stimuli, however only if the respective SPN was still active when the RPE was signalled at US by VTA DAN (***Oettl et al., 2020***). We go beyond this work by formally connecting representation changes to temporal difference learning to predict how individual SPN responses should change at a trial-by-trial level.

Besides sensory stimuli, a critical aspect of environmental states that predicts reward in natural settings and Pavlovian conditioning paradigms is the representation of time. Prior theoretical work has modeled dopamine’s role in setting an internal clock that keeps subjective time (***Mikhael and Gershman, 2019; Jakob et al., 2022***). In this model, an upstream time representation (e.g., from striatal neurons; ***Mello et al., 2015***) is transformed into subjective time by a single scaling parameter. A linear value function approximation is used, and the subjective time parameter is learned by minimizing reward prediction error, similar to our set-up. The authors confirm a variety of predictions of the model in both behavioral and dopamine recording data.

We go beyond these prior works in two respects. First, we analyze the stimulus representation in conjunction with the RPE signaled by dopamine and provide direct evidence for trial-by-trial error-driven updates to stimulus representations underlying value estimation. Second, our updates are distinct for each neuron’s parameters, as opposed to a single global parameter, providing a more detailed prescription for representational changes. Thus, we propose that appropriate simultaneous recordings of dopamine and subjective time-coding striatal projection neurons could be used to test whether time representation learning also occurs by RPE minimization using the methodology presented in this work.

Beyond the OTu, midbrain dopamine neuron subpopulations project to targets including the nucleus accumbens, dorsal striatum, and cortex (especially medial prefrontal cortex). There is evidence the nucleus accumbens also represents states that support value estimation. A natural extension of our OTu representation learning hypothesis is that the structure of nucleus accumbens representations also depends on dopamine error-driven learning since it shares cell types and regional connectivity. Like OTu, nucleus accumbens is thought to encode state or value information upstream of VTA (***Winkelmeier et al., 2022***). Additionally, it has recently been shown that D1-expressing SPNs and DANs in the nucleus accumbens-VTA circuit are hardwired to compute a temporal-difference error if SPN response magnitude corresponds to value (***Campbell et al., 2025***). Thus, we hypothesize that dual recordings in nucleus accumbens and VTA during Pavlovian conditioning would show representation changes consistent with the RPE minimization prescription.

### Latent weight inference

To make predictions using the gradient descent update, we needed the SPN output weight, which was not experimentally available. We developed a new method for weight inference by casting the gradient descent update to **w** as a linear-Gaussian dynamical system with **w** as latent variables.

Inference for linear-Gaussian models is analytically tractable using Kalman filtering and smoothing and yields exact posterior mean and covariance parameters for the trajectory of **w** values. Beyond our particular use case, the method is more generally applicable to inferring the trial-by-trial time-series of synaptic connectivity between groups of recorded neurons. The weight dynamics can be modified (e.g., to Hebbian plasticity) as long as the dynamics are linear and the process noise model is Gaussian. In the observation model, **s** can be replaced with the upstream neural population activity, and *δ* can be replaced with the downstream neural population activity. The form of observations would need to be chosen to reflect the neurobiology and could incorporate nonlinear activations, as long as it is safe to assume Gaussian additive noise to capture additional variability.

### Limitations and future directions

A limitation of our current analysis is that we cannot disambiguate stimulus and state representations in the OTu. In the experiment we analyze, the task states are specified by the odor presented during the stimulus period, which differentially predict reward probability. However, in general, it is possible that multiple configurations of sensory inputs could correspond to the same task state. For example, an animal could learn that odors *A* and *B* both (independently) predict 100% probability of reward. Then, after a contingency change where *A* and *B* now predict 0% reward probability and the animal experiences some trials with odor *A* and no trials with odor *B*, what should happen to the representation of odor *B*? If the OTu represented task state, then we would predict the response to *B* changes in a way that is coupled to how the response to *A* changes. Alternatively, the representation of *B* may not change if OTu SPNs only represent stimuli and not task states. There is suggestive evidence that stimuli are bound to state representations in OTu because D1-receptor expressing SPNs quickly learn to represent odors based on the associated reward (which is the relevant distinction between states), and representations of same-valence odors evolve similarly in neural state space (***Martiros et al., 2022; Millman and Murthy, 2020***). Future experiments that map multiple stimuli to the same state or require the animal to infer the state in a probabilistic paradigm may be used to confirm that SPN response changes correspond to state representation updates by dopamine signals.

In terms of the learning algorithm, we acknowledge that the backpropagation of exact weight information in the gradient descent rule is biologically implausible. However, ventral striatum SPNs typically exclusively express D1 or D2-receptors, imposing specific third factor Hebbian plasticity rules (***Pawlak and Kerr, 2008; Shen et al., 2008; Sosis and Rubin, 2025***). Thus, the outgoing synaptic weight’s sign information is plausibly available at the input synapse, making the update local. Consistent with this, we found that the predictivity of the gradient descent update is mostly preserved if **w** is replaced with the sign of **w** in Figure 4. This result is also coherent with a prior computational study that relaxes the exact weight transport problem by backpropagating correct weight signs with random feedback weight magnitude to achieve similar performance to stochastic gradient descent when training neural networks (***Liao et al., 2015***).

Methodologically, we did not constrain the sign of **w** when using the smoother for inference. This sometimes resulted in **w** sign changes, which may be biologically unrealistic, since expression of D1- or D2-receptors imposes specific plasticity rules that remain to be fully characterized (***Pawlak and Kerr, 2008; Sosis and Rubin, 2025***). It is possible to re-formulate the weight inference problem to constrain the sign by using a hidden Markov model or by giving up analytic tractability and using a posterior sampling-based approach. The former approach has the drawback of discretizing hidden states somewhat arbitrarily, while the latter is more computationally expensive and often requires carefully tuning the inference algorithm. Moreover, effective sign changes may be implemented between SPNs and DANs by interspersed GABA neurons in the VTA and ventral pallidum. Thus, we preferred the relatively simple and efficient smoother approach. An interesting future direction would be to record from identified D1/D2-receptor expressing SPNs in conjunction with DANs. This would enable validation of the weight inference method and testing whether definitively knowing the sign of the linear approximation weight improves representation update predictivity.

Lastly, our choices for stimulus encoding and state representation are debatable. The olfactory bulb and piriform cortex have sparse odor representations and support excellent identity decoding, so the one-hot encoding simplification may be a good approximation of 3 nearly orthogonal responses to unrelated cue odors. However, if multiple odors need to be mapped to the same state and updated together, then they must become coupled in some way as supported by recent work on OTu SPNs (***Millman and Murthy, 2020***). In the present analysis, with one odor per state, this issue is not relevant. However, to handle multiple odors corresponding to a state, a different parameterization is needed.

### Concluding remarks

In summary, our analyses reveal that changes in OTu SPN responses to odor stimuli during Pavlovian conditioning are consistent with gradient descent on RPE signaled by dopamine. By comparing to several alternative updates, we show the RPE and sign of output weights are essential to predictivity. These results demonstrate error-driven representation learning in the brain at a trial-by-trial level. Our study provides initial support for the broader hypothesis that dopamine signals throughout the brain are a substrate for error-driven representation learning.

## Acknowledgments

We are grateful to Farhad Pashakhanloo for helpful discussions. This work was supported by the Kempner Institute for the Study of Natural and Artificial Intelligence (G.C., S.J.G.), the Department of Defense MURI program under ARO grant W911NF-2310277 (S.J.G.), DFG grant KE1661/9-1 (W.K.).

## Methods

### Experimental overview

All procedures were in accordance with the National Institutes of Health Guide for the Care and Use of Laboratory Animals and the EU 2010/63 directive, and approved by the local animal welfare authority (Referat 35, Regierungspräsidium Karlsruhe, Karlsruhe, Germany). Adult male mice were trained head-fixed on an olfactory Go/No-Go task. Trials comprised a variable ITI (8–12 s) with clean air, a 1 s odor cue, and a 1 s post-cue response window during which anticipatory licking was scored. Water reward was delivered at the end of the response window. Access to water was controlled for the mice in order to maintain their motivation to obtain rewards during the task.

Training proceeded through three stages: (i) shaping, in which a single odor predicted certain reward, (ii) discrimination, in which three odors were presented pseudorandomly with reward probabilities 100%, 50%, and 0% (100% and 50% were “Go”; 0% was “No-Go”), and (iii) reversal, in which the 100% and 0% odor contingencies were swapped after stable performance. Performance was quantified as the fraction of correct trials (Hits on Go trials and Correct Rejections on No-Go trials).

Subjects were adult DAT:(IRES)Cre^+^ mice (B6.SJL-Slc6a3tm1.1(cre)Bkmn/J; RRID: IMSR_JAX:006660; Jackson Laboratories) crossed to a Cre-dependent ChR2 reporter line (Ai32(RCL-ChR2(H134R)/EYFP, Jackson Laboratories), enabling optogenetic tagging of dopamine neurons; backcrossed >F10 in a C57BL/6J background (Charles River)). Eight animals were chronically implanted with custom dual-region tetrode arrays and optical fibers for simultaneous recordings from olfactory tubercle (Tu) and ventral tegmental area (VTA).

Behavior and neural data were acquired and synchronized with task events (odor, licking, reward, laser) and respiration (pressure sensor). Spikes were sorted with Kilosort and curated using automated criteria for spatial localization, refractory-period violations, lick-artifact contamination, and redundant cluster merging.

Putative VTA dopamine neurons were identified via optogenetic tagging using spike-laser cross-correlograms to assess short-latency activation. Striatal projection neurons (SPNs) in Tu were operationally defined as recorded units with ITI firing rates below 5 Hz at the end of each session. The full experimental methods are available in (***Scheller et al., 2026***).

### Analysis criteria and preprocessing

For the main analyses in this paper, we required a minimum of 8 SPNs and 8 DANs present in the session. To normalize SPN and DAN firing rates, we consider the window [-2.5 s,-0.5 s] relative to stimulus onset for each trial. We subtract these intertrial pre-CS spike rates from the stimulus period [0 s, 1 s] and reward period [2 s, 2.5 s] spike rates to compute SPN and DAN responses. We use the intertrial interval for normalization due to ramping in the DAN firing rates before the reward period, although we consider alternative thresholds for neuron counts and normalization windows described in the supplement. No additional normalization is performed.

### Neural response predictions

To map the model to the experiment, we have measurements in each trial of reward *r*, RPE *δ* via average normalized DAN activity during the reward period, and **s** via the population activity of SPNs during the stimulus period. We infer **w** and *v* to be consistent with *r* and *δ* at each trial (described below in Latent weight trajectory inference). Assuming we have 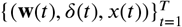 for the entire session of *T* trials, the update is Δ*θ*_*k*_(*t*) ∝ *δ*(*t*)**w**(*t*) for odor *k*. In the data, we compute Δ**s**_*k*_(*t*) = **s**_*k*_(*t*^′^) − **s**_*k*_(*t*) where **s**_*k*_(*t*) is the response to odor *k* at trial *t*, and **s**_*k*_(*t*^′^) is the SPN population response in the next trial where odor *k* is presented. To measure predictivity, we compute the Pearson correlation of the two update time series for each neuron. We compute dependent *t*-tests for paired samples with a one-sided alternative that the rule has greater mean correlation, treating each neuron as a paired sample.

We additionally group all predicted updates for all SPNs and trials by decile and plot these deciles against the corresponding changes in SPN response to an odor across successive presentations. In total, 99902 updates were analyzed, resulting in approximately 9990 updates per decile to average over.

### Latent weight trajectory inference

The weight update in Eq. 4 is a linear dynamical system. We treat the weights as latent variables and RPE *δ* and reward *r* as observed variables at each time step, as shown in Figure 1**B**. If we assume additive Gaussian noise in the weight updates and observations, then this is a linear-Gaussian system. Specifically,

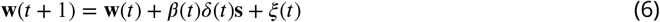

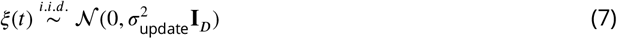

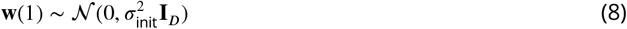

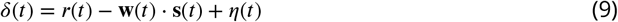

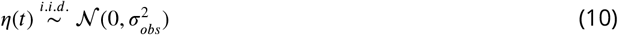

where the noise terms, *ξ*(*t*) and *η*(*t*), are sampled independently at each step *t* = 1, …, *T* − 1. We make the simplifying assumption that the noise is isotropic and the initial weights are multivariate Gaussian. We make an additional assumption that the subjective value of the water reward is constant throughout the session even as the mouse reaches satiety. Lastly, we parameterize the learning rate as 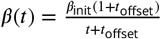, which permits delaying the learning rate decay through *t*_offset_ and scales the learning rate through *β*_init_. In the case where learning rate is held constant, we simply set and infer *β*(*t*) = *β*_init_.

The Rauch-Tung-Striebel smoother infers the posterior probability of latent weights given the full sequence of observations Pr(**w**(*t*) | *δ*(1), …, *δ*(*T*)). The posterior mean provides the most probable value of **w**(*t*), which we use in the gradient update to SPN representation. The process also depends on hyperparameters, which need to be inferred by an outer loop of maximizing the marginal likelihood of the observations. We place a weak log-normal prior on *σ*_init_ with (*μ* = log(0.5), *σ* = 1). We use `scipy.optimize.minimize` on the negative log-likelihood. We list the hyperparameters, bounds, and initial values in Table 1.

**Table 1.** Hyperparameter bounds and initial values for marginal likelihood optimization procedure.

| parameter | bounds | initial value |
| --- | --- | --- |
| $\sigma_{\text{update}}$ | $(10^{-8}, 0.001)$ | 0.001 |
| $\sigma_{\text{obs}}$ | $(10^{-8}, 0.001)$ | 0.001 |
| $\sigma_{\text{init}}$ | $(10^{-2}, 5.0)$ | 1.0 |
| $\beta_{\text{init}}$ | $(5 \cdot 10^{-4}, 5.0)$ | 0.05 |
| $t_{\text{offset}}$ | $(0, 10)$ | 0.0 |

### Model recovery from synthetic data

Using the exact sequence of odors, rewards, and DAN responses in the experiment, we simulate synthetic trajectories of **w** and **s** according to the latent weight dynamics and representation update models, respectively. We use hyperparameters inferred from the experimental data for each session to generate the synthetic data in the same regime as the experiment. Then, the exact same weight recovery and hyperparameter optimization procedure are used again.

Because a multi-dimensional latent variable is mapped to a scalar observation in the smoother, and our prior on the weights is symmetric, there can be a degeneracy in solution modes. As such, we heuristically require the posterior weight variance to decrease by 10% relative to the prior weight variance in 10% of trials to consider a weight identified. Unidentified weights and weights that were below a threshold magnitude of 10^−5^ were omitted in the downstream recovery analysis. All weights were identifiable in the real data, but this was not the case in the synthetic data.

For a posterior predictive check, we sample 1000 weight trajectories from the posterior and compare the samples to the synthetic weight trajectory by Pearson correlation (Figure 2**A**). Then, we produce sampled trajectory updates to the SPN representation according to eq. 4 with 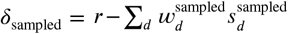, and compute Pearson correlation with the synthetic updates (Figure 2**B**). Lastly, to compute the update prediction correlation, we plug the weight trajectory means and synthetically generated *δ* trajectory into eq. 4 and compare to the changes in the synthetically generated **s** (Figure 2**C**).

### Gradient descent ablations and alternative models

To ablate the contribution of RPE, we fix *δ*(*t*) = 1 in the latent weight update for all trials while retaining the DAN activity *δ*(*t*) in the observations. We re-fit linear-Gaussian system parameters. We use the Rauch-Tung-Striebel smoother to infer weights accordingly. Using these inferred weights and *δ*(*t*) = 1, we compute the update correlations as eq. 4. To ablate the effect of output weights for credit assignment, we set **w**(*t*) = **1** when computing SPN updates. To test if a local version of the rule based on output weight sign predicts the update, we replace **w** with sign(**w**) element-wise. For random feedback alignment, we replaced the trajectory of **w** with a single fixed weight vector **w**^*F A*^ for all trials. The vector was inferred by setting the learning rate and process noise to 0 (*β*_*init*_ = 0, *σ*_*update*_ = 0) in the smoother, fitting the remaining parameters, and inferring the weight posterior. We compute the predicted update and correlation to the observed update using the fixed inferred weight **w**^*F A*^. Lastly, in the random sign model, we randomly sample a sequence of RandomSign(0.5) variables and modify the linear-Gaussian dynamics with these random signs when fitting parameters. Then, we use the Kalman smoother to infer a posterior over weight trajectories and compute updates using these weights.

### Held out comparison for learning rules

We used the first 120/150 trials (80%) in each session as a training set to optimize the hyperparameters of the Kalman smoother for each update rule as described in the previous section. Then, we run the Kalman smoother on all trials to infer the full weight trajectory. The correlation between the predicted updates of the last 30 trials (20%) and measured response changes are reported in the held out comparison figures. Updates were considered part of the last 30 trials if the second time point of the pair was in the last 30 trials.

### Bayesian information criterion comparison

To compare models’ ability to explain the data, we compute the BIC using the marginal likelihood provided by the Kalman smoother after hyperparameter optimization using

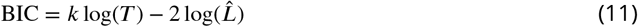

where *k* was the number of hyperparameters, *T* = 150 was the number of trials, and 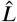 was the maximized marginal log-likelihood (***Bishop, 2006***). The sign(**w**) rule is not available in the BIC comparison because we did not have an analytic expression for the likelihood associated with the weight sign.

## Supplementary Figures

**Figure S1.**
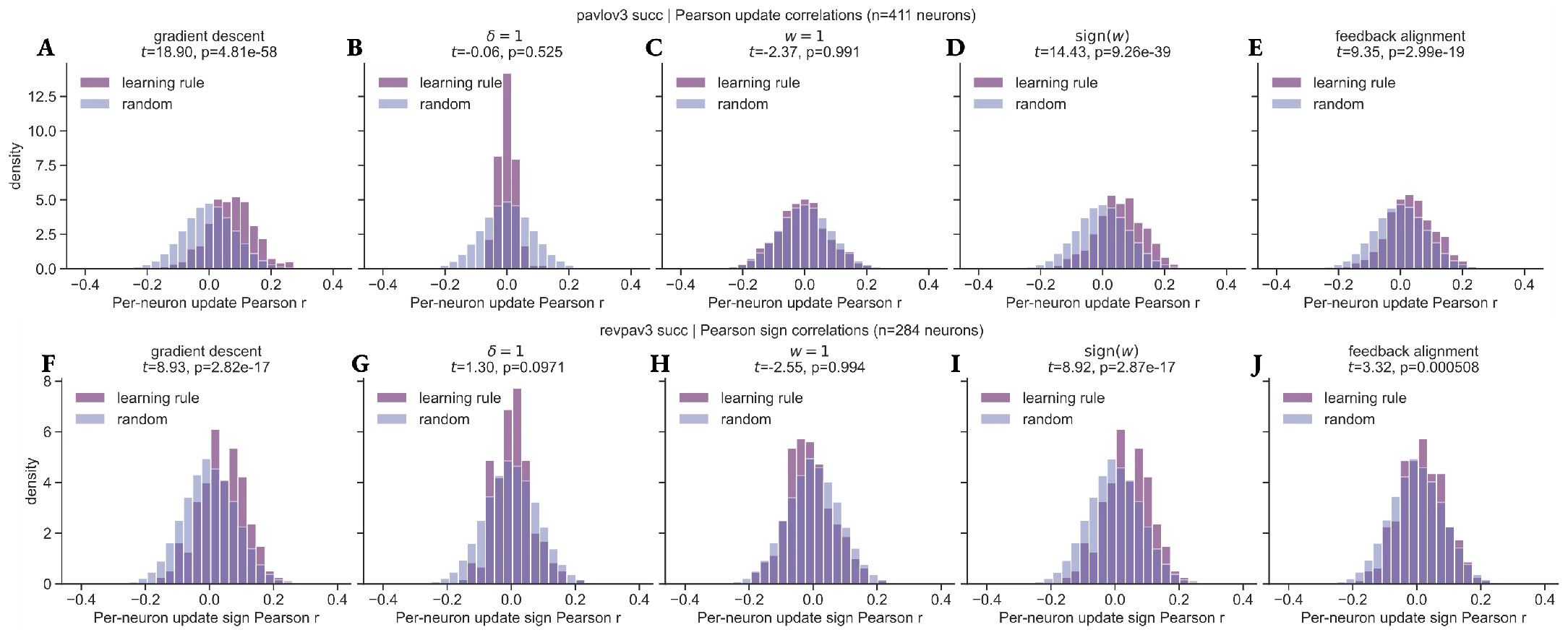
Full histogram of correlations between successive model-inferred and recorded changes in SPN response. Top row: Pavlovian conditioning sessions (411 neurons). Bottom row: Reversal sessions (284 neurons). Our alternative representation updates are: (**A, F**) Gradient descent rule, (**B, G**) fixing *δ* = 1, (**C, H**) fixing **w** = **1**, (**D, I**) replacing **w** by its element-wise sign, (**E, J**) replacing **w** by a fixed Gaussian vector. Results of paired *t*-tests against a random-sign shuffle null of each model’s updates are shown above each plot.

## Data and code availability

Data will become publicly available upon the publication of (***Scheller et al., 2026***). Code will become publicly available on https://github.com/george-tog/DA_rep_learning upon review.

**Figure S2.**
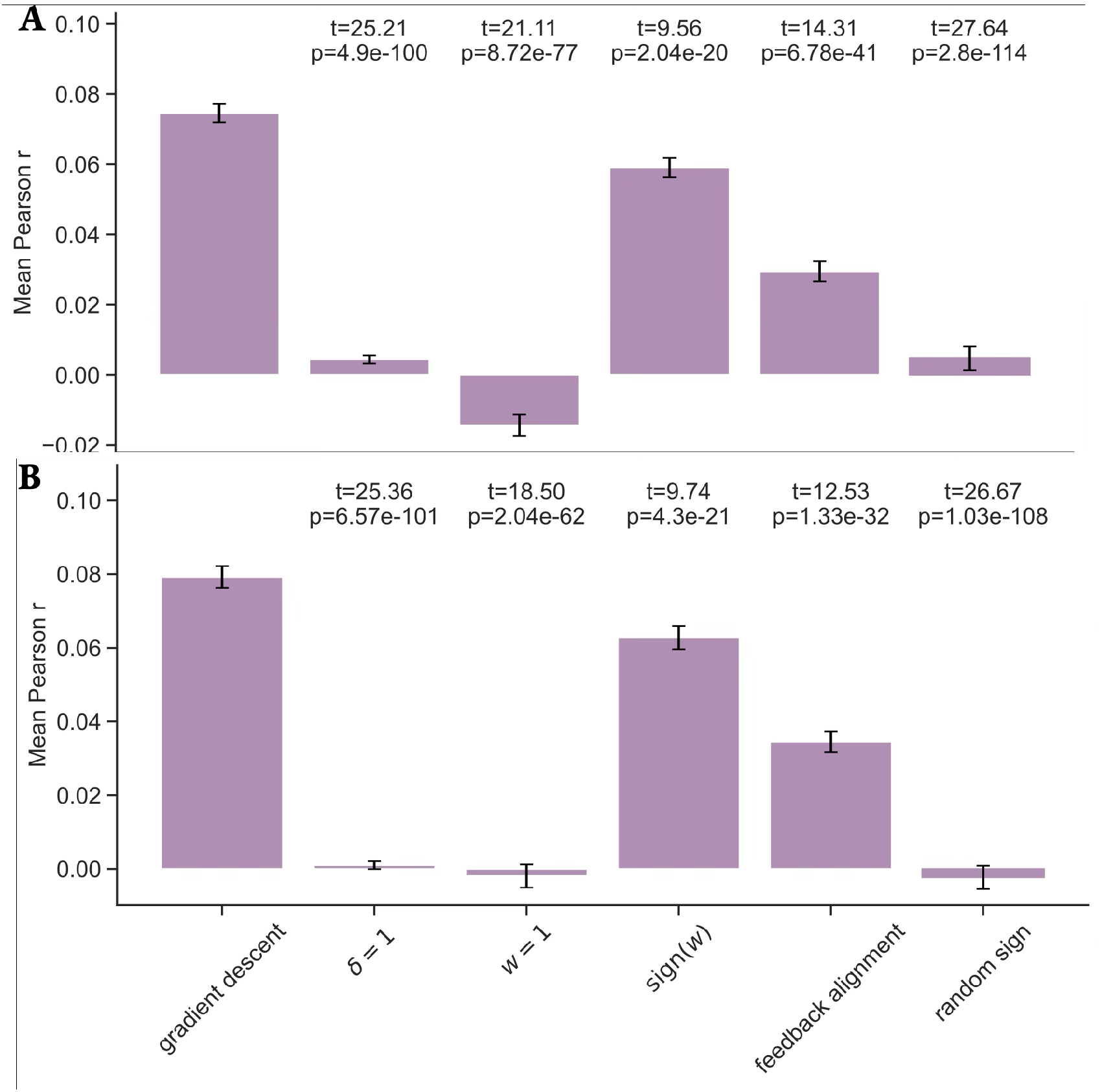
Normalization window robustness of update correlation results across both Pavlovian conditioning and reversal sessions. (**A**) 2 s pre-CS window, 0.5 s offset from stimulus and response window for SPNs and DANs respectively. (**B**) 1 s pre-CS window, 0.5 s offset from stimulus window for both SPNs and DANs. Bars show mean and standard error across neurons.

**Figure S3.**
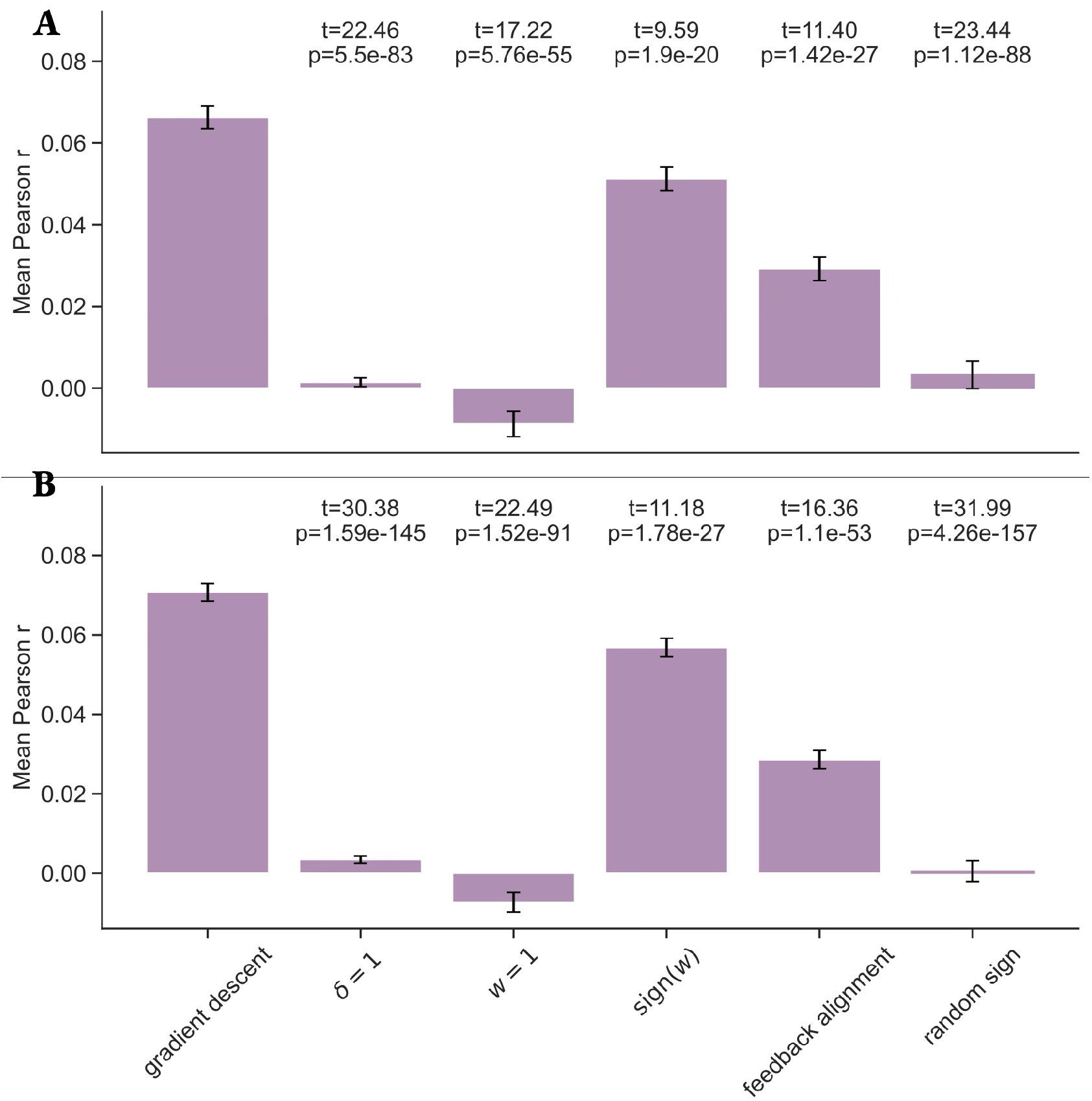
Robustness to minimal SPN and DAN count thresholds. Results are robust to requiring at least (**A**) 16 SPNs and 8 DANs and (**B**) 8 SPNs and 4 DANs per session. Bars show mean and standard error across neurons.

**Figure S4.**
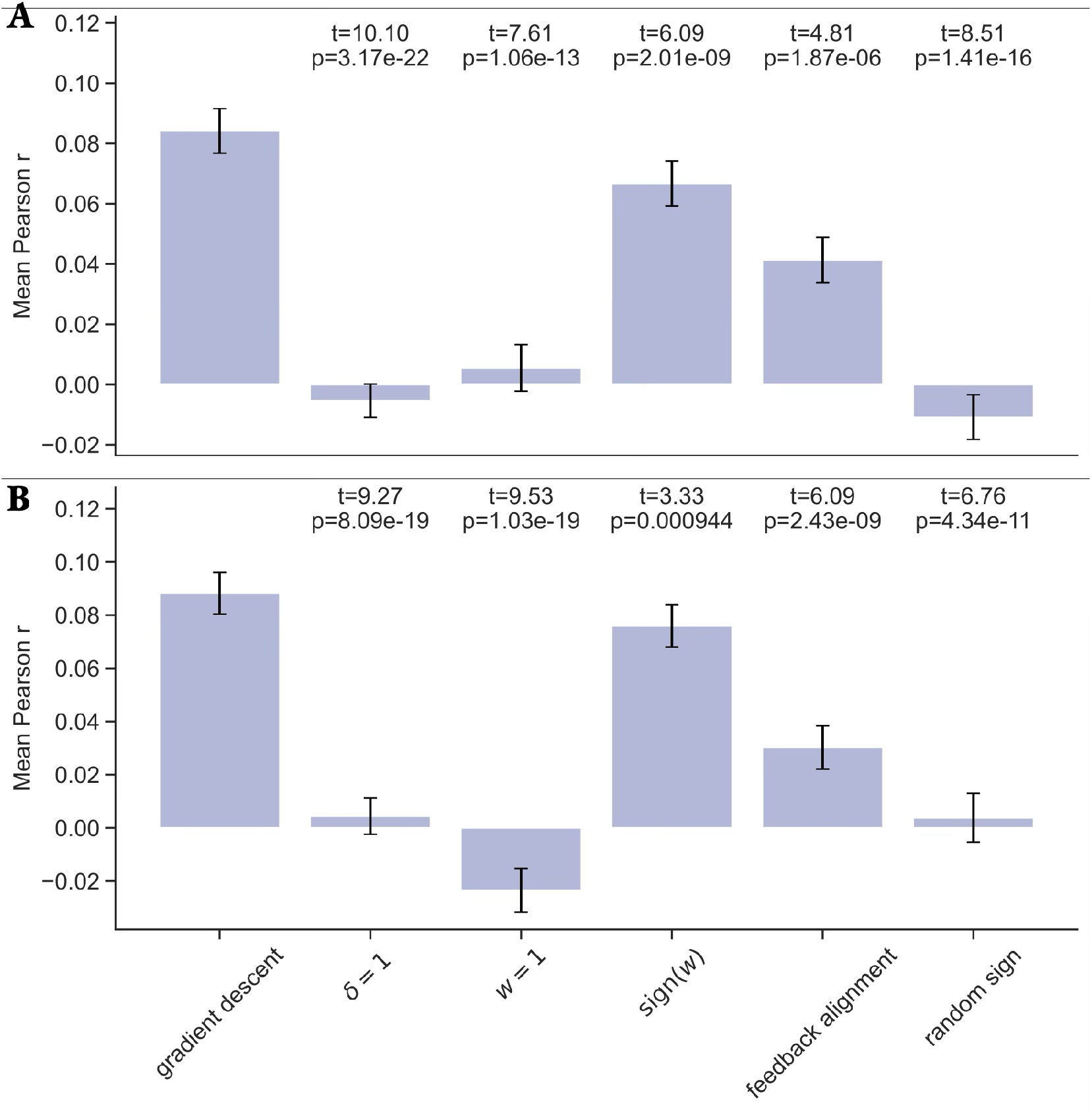
Correlations between predicted and observed changes in olfactory tubercle activity pooled across (**A**) Pavlovian conditioning sessions and (**B**) reversal sessions when predictions are evaluated on data held out with respect to weight inference hyperparameter fitting. Bars show mean and standard error across neurons. The gradient model (Eq. 5) was compared to several alternative representation updates: fixing *δ* = 1, fixing **w** = **1**, replacing **w** by its element-wise sign (sign(**w**)), replacing **w** by a fixed Gaussian vector (feedback alignment), and randomly sampling the sign of the update (null). The *t*-values and *p*-values shown above each alternative model are derived from paired *t*-tests. Bars show mean and standard error across neurons.

**Figure S5.**
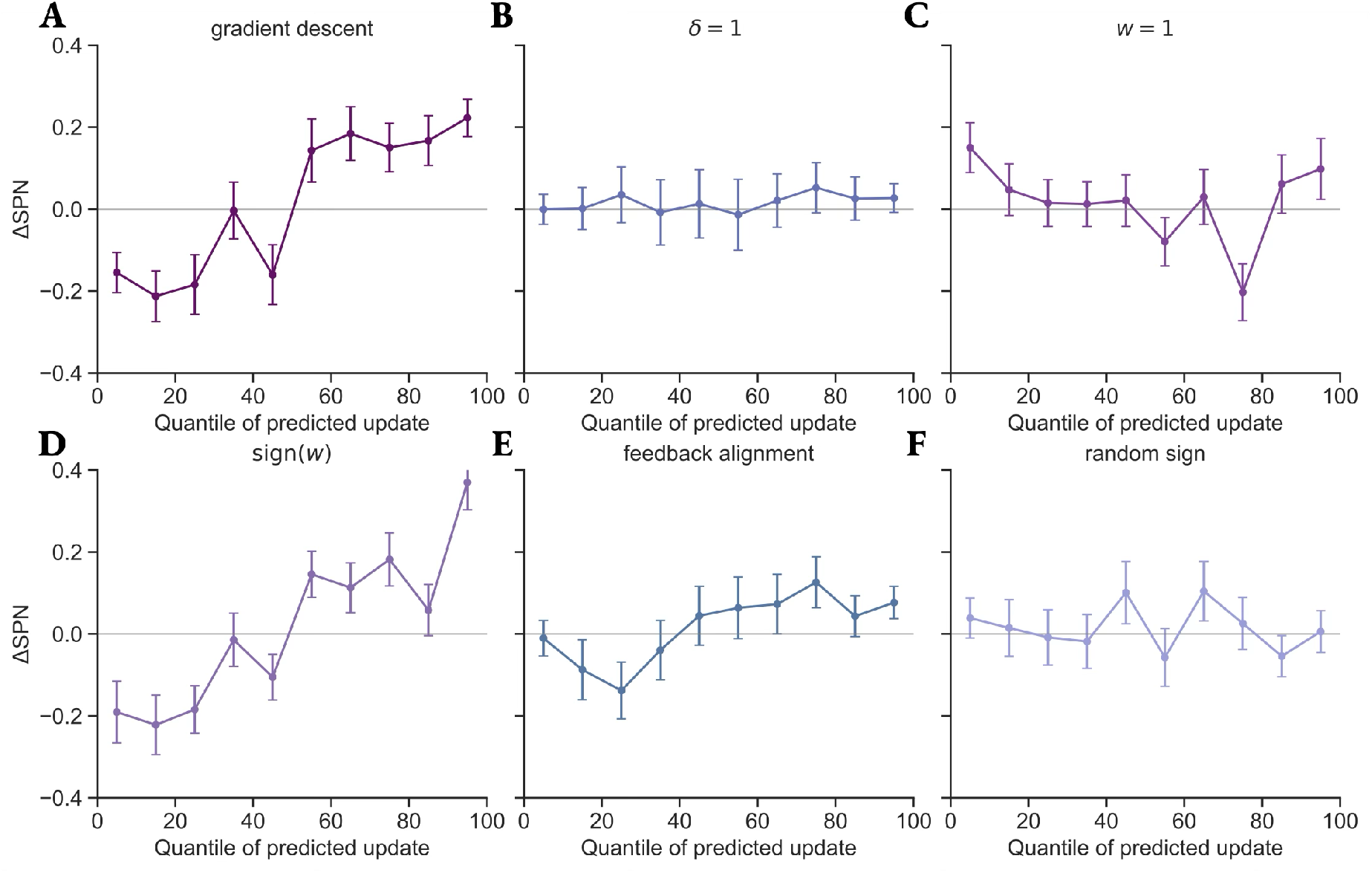
Predicted SPN changes computed on trials that were held out from hyperparameter fitting in the weight inference procedure. Change in SPN response between successive representations of an odor as a function of the decile of predicted change according to (**A**) gradient descent, (**B**) *δ* = 1, (**C**) **w** = **1**, (**D**) sign(**w**), (**E**) feedback alignment, (**F**) random sign update rules. Points denote mean and error bars denote standard error. A total of *n* = 20443 updates were considered across all SPNs and trials.

**Figure S6.**
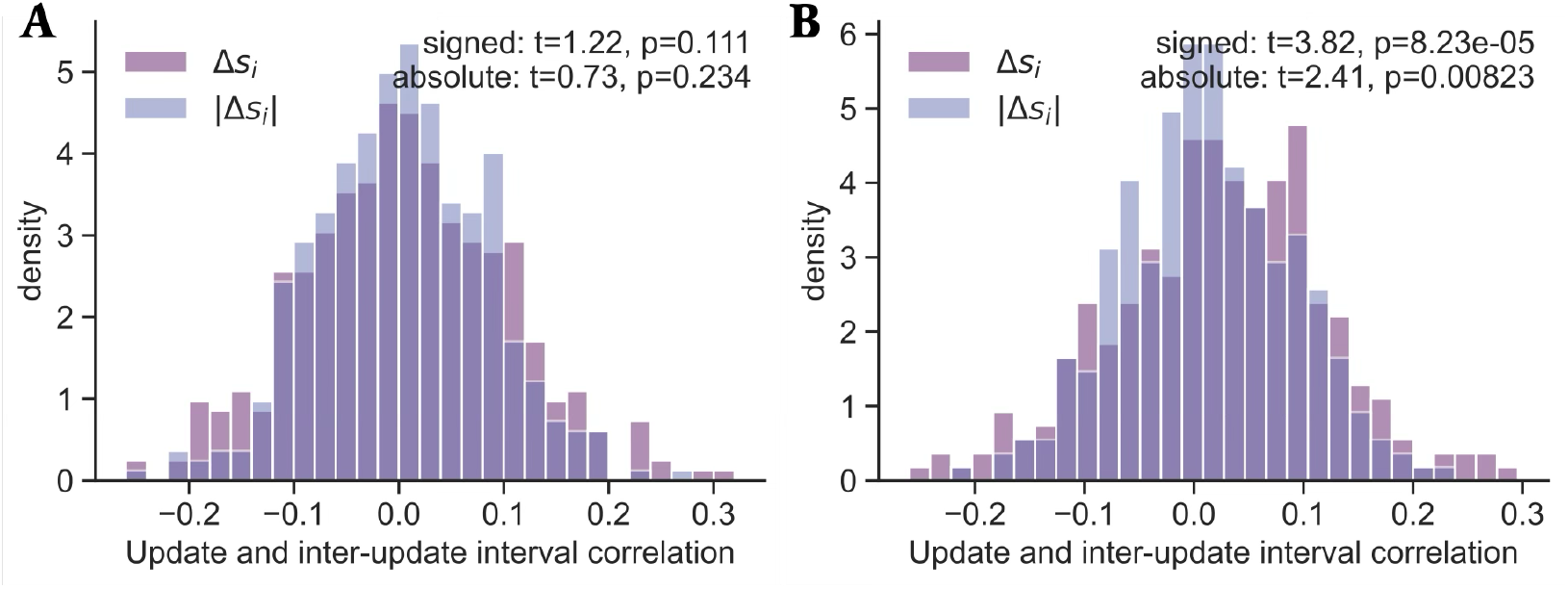
Correlation between each SPN update and the number of trials between successive measurements of SPN response to a particular odor stimulus. Pavlovian conditioning (**A**) and reversal (**B**) sessions are shown. We compute a 1-sample *t*-test against the alternative hypothesis that the mean is greater than 0.

**Figure S7.**
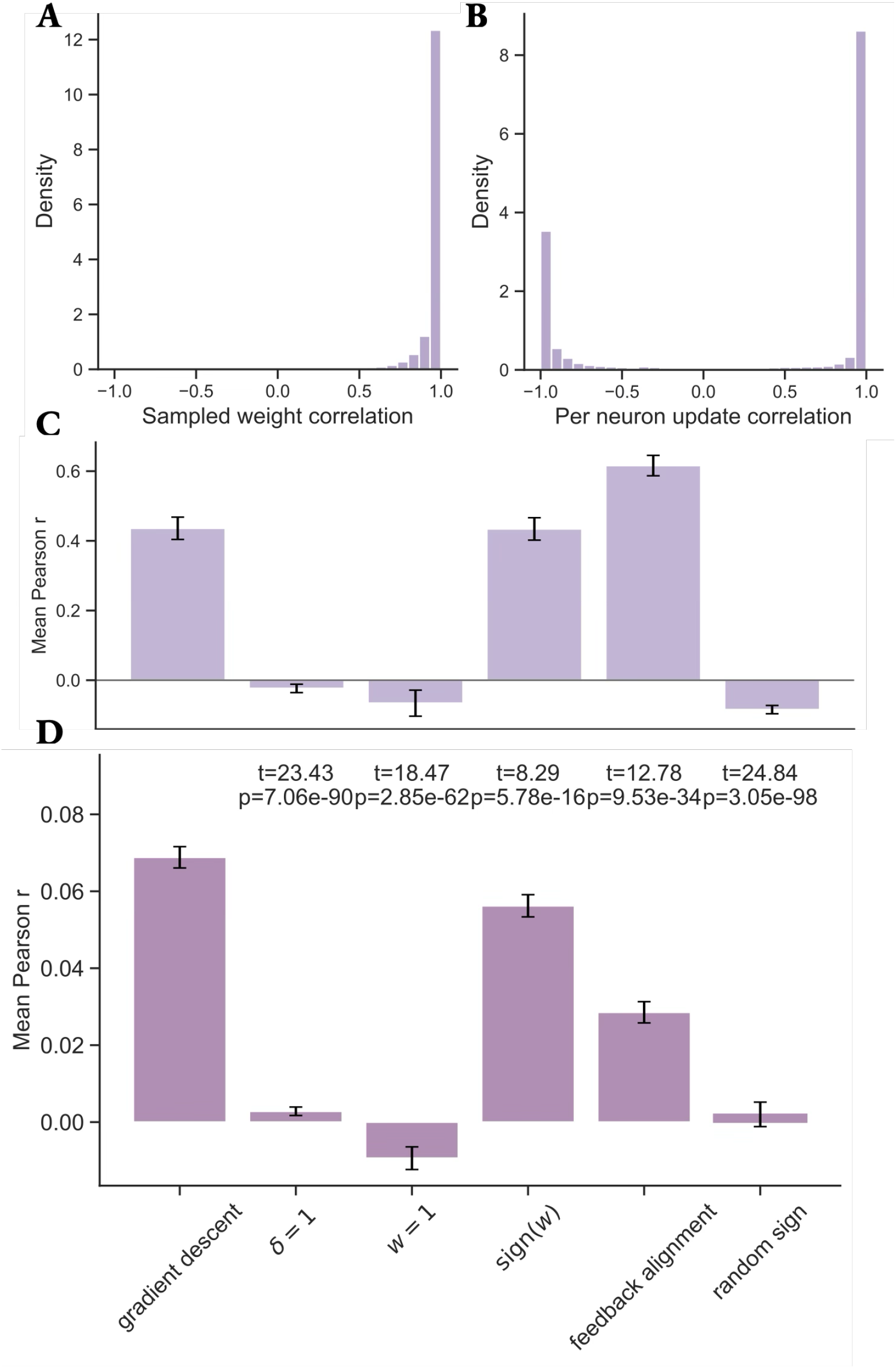
Weight inference synthetic recovery (**A**-**C**) and update predictions (**D**) with inferred constant weight learning rate *β*. (**A**) Correlation between the true and inferred weight trajectory. (**B**) Correlation between true and reconstructed representation updates from sampling weight trajectories and reconstructing RPEs and representation trajectories. **C** Correlation between true and inferred representation updates using weight trajectory mean and synthetic RPEs. (**D**) Predicted update correlation on the data. Results are aggregated across both Pavlovian conditioning and reversal sessions. (**C**-**D**) Bars show mean and standard error across neurons.

